# A 2.8 Å structure of zoliflodacin in a DNA-cleavage complex with *Staphylococcus aureus* DNA gyrase

**DOI:** 10.1101/2022.11.30.518515

**Authors:** Harry Morgan, Magdalena Lipka-Lloyd, Anna J. Warren, Naomi Hughes, John Holmes, Nicolas P. Burton, Eshwar Mahenthiralingam, Ben D. Bax

## Abstract

Since 2000 some thirteen quinolones/fluoroquinolones have been developed and come to market. The quinolones, one of the most successful classes of antibacterial drugs, stabilize DNA-cleavage complexes with DNA gyrase and topo IV, the two bacterial type IIA topoisomerases. The dual targeting of gyrase and topo IV helps decrease the likelihood of resistance developing. Here we report a 2.8 Å X-ray crystal structure which shows that zoliflodacin, a spiropyrimidinetrione antibiotic, binds in the same DNA-cleavage site(s) as quinolones sterically blocking DNA religation. The structure shows that zoliflodacin interacts with highly conserved residues on GyrB (and does not use the quinolone water-metal ion bridge to GyrA) suggesting it may be more difficult for bacteria to develop target mediated resistance. We found that zoliflodacin had an MIC of 4 *µ*g/*m*L against *Acinetobacter baumannii*, an improvement of 4-fold over its progenitor QPT-1. The current phase III clinical trial of zoliflodacin for gonorrhea is due to be read out in 2023. Zoliflodacin, together with the unrelated novel bacterial topoisomerase inhibitor gepotidacin, are likely to become the first entirely novel chemical entities approved against Gram-negative bacteria in the 21st century. Zoliflodacin may also become the progenitor of a new safer class of antibacterial drugs against other problematic Gram-negative bacteria.

## 1. Introduction

Zoliflodacin is an oral spiropyrimidinetrione antibiotic currently in a phase III clinical trial for the treatment of gonorrhea, a sexually transmitted infection (STI) caused by the Gram-negative bacteria *Neisseria gonorrhoeae* [1-4]. Zoliflodacin (Figure 1a) was developed from QPT-1 (or PNU-286607) a compound discovered in Pharmacia by whole cell screening against Gram-negative (and Gram-positive) bacteria [5]. QPT-1 (Figure 1b), discovered for its antibacterial whole cell activity, was found to inhibit the bacterial type IIA topoisomerases; *E. coli* DNA gyrase (IC_50_ 9 *µ*M) and *E. coli* topoisomerase IV (IC_50_ 30 *µ*M) [5]. This method of discovery is reminiscent of the discovery of quinolone/fluoroquinolone antibiotics, also initially discovered for whole cell activity and then found to be inhibitors of the bacterial type IIA topoisomerases [6].

**Figure 1.**
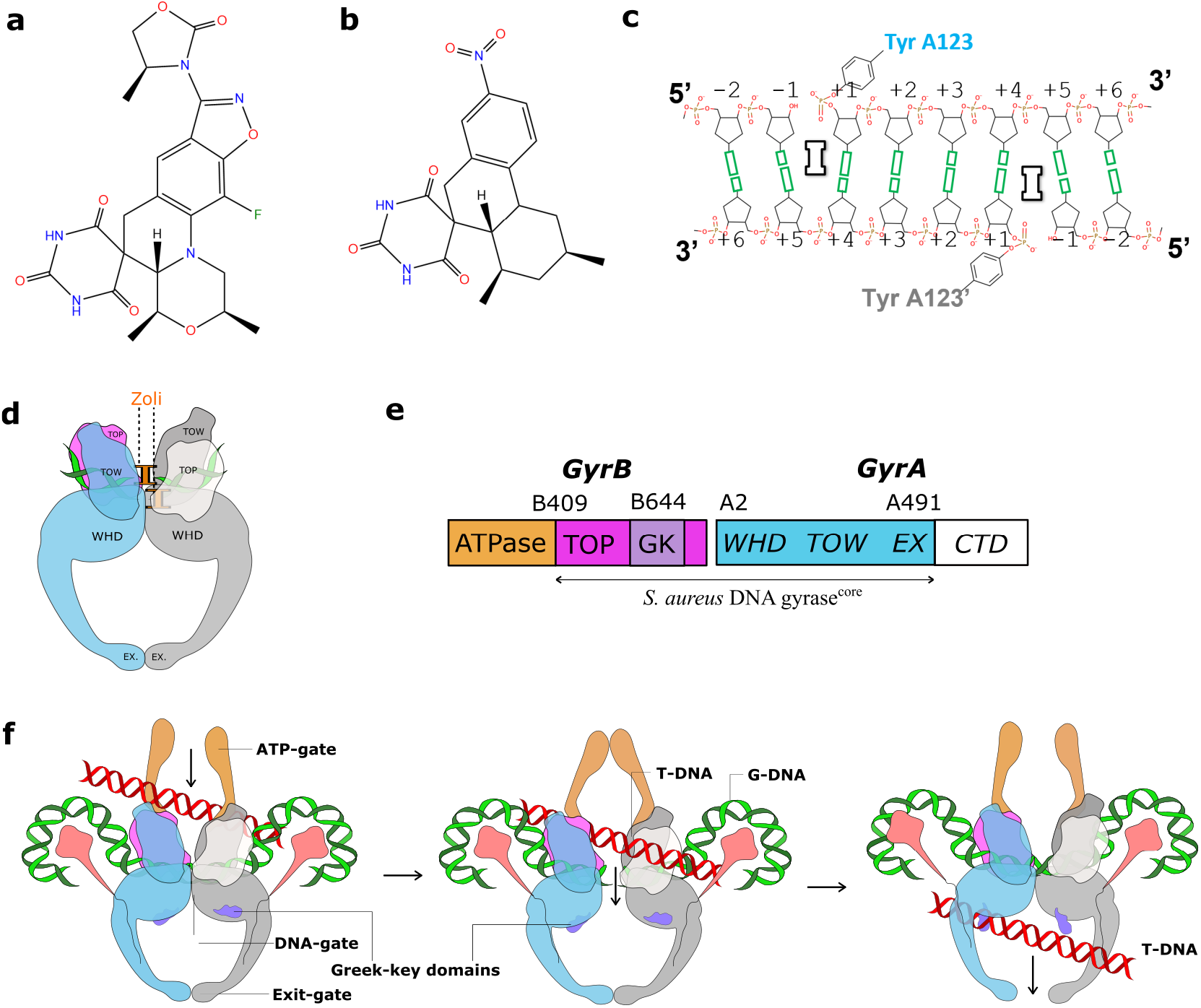
Zoliflodacin, QPT-1 and *S. aureus* DNA gyrase DNA-cleavage complexes. (**a**) Chemical structure of zoliflodacin (oxygens red, nitrogens blue, fluorine green) (**b**) chemical structure of QPT-1 (**c**) A schematic of the central eight base-pairs of DNA, with two inhibitors (I) binding in the cleaved DNA and inhibiting DNA religation. Note that DNA-cleavage takes place between the -1 and +1 nucleotides on both the Watson and Crick strands (**d**) A schematic of the DNA-cleavage complex with two zoliflodacins *S. aureus* DNA gyrase and DNA presented in this paper. (**e**) The *S. aureus* DNA gyraseCORE construct used consists of residues B409-B644 from GyrB, fused to A2 to A491 from GyrA. The small greek key (GK) domain has been deleted from GyrB [13] (**f**) A simplified schematic of DNA gyrase, in which a G-DNA duplex (green) is cleaved by the enzyme and another DNA duplex (known as the T or transported DNA - red) is moved through the enzyme. The greek key domains are not involved in cleaving the gate (or G) -DNA segment [10, 13]. The C-terminal domains (CTD) are shown in pink (approx. positions as in full length *E. coli* structures [14]).

Topoisomerases are essential enzymes needed to relieve topological problems when the DNA double helix is unwound for both DNA replication and transcription [7]. Topoisomerases are divided into type I topoisomerases, that introduce single-stranded DNA breaks to modify DNA topology and type II topoisomerases that modify topology by introducing double-stranded DNA breaks [7-9]. Most bacteria possess two type IIA topoisomerases, DNA gyrase and topo IV. While DNA gyrase can uniquely introduce negative supercoils into DNA, topo IV has good decatenase activity [9, 10]. A mechanism for topological changes introduced by DNA gyrase is shown in Figure 1.

The introduction of double-stranded breaks into DNA is potentially hazardous for the cell and the stabilization of DNA-cleavage complexes by quinolones is often bactericidal [11, 12].

The proposal that Gram-negative bacteria evolved a second cell wall to protect them from antibiotics produced by other micro-organisms [15] may partly explain the failure of new classes of antibiotics targeting Gram-negative bacteria, to date, in the 21st century [16-18]. Perhaps for Gram-negative bacteria the hardest task is to get into the cells and a whole cell screening approach followed by target identification of proven targets is more likely to be successful [19, 20]. Indeed, GlaxoSmithKline discovered and developed the NBTI gepotidacin, another new class of DNA gyrase targeting antibiotic currently in phase III clinical trials [21], from a hit compound active in a screen for whole cell antibacterial activity [13]. The chemical diversity of NBTIs such as gepotidacin, which stabilize single stranded DNA-cleavage complexes with bacterial type IIA topoisomerases, suggested that this class of compounds could not have a chemistry-based name [13, 22-27]. The name NBTI, although originally a pneumonic for <u>n</u>ovel <u>b</u>acterial topoisomerase inhibitor [13], could also be taken to stand for <u>N</u>on-DNA-cleavage pocket <u>B</u>inding on the <u>T</u>wofold-axis Inhibitor (as this describes the binding mode of the chemically diverse NBTIs [13, 22-27]).

The occurrence of antimicrobial resistance in hospital acquired ESKAPE pathogens (*Enterococcus faecium, Staphylococcus aureus, Klebsiella pneumoniae, Acinetobacter baumannii, Pseudomonas aeruginosa*, and *Enterobacter* species) was a major cause for concern in 2009 [28]. New classes of antibiotics have now been developed for Gram-positive bacteria, such as the tiacumicin Fidaxomicin for *Clostridioides difficile* [29]. However, Gram-negative bacteria (the KAPE in ESKAPE) remain a major cause for concern. The popular quinolone/fluoroquinolone class of antibacterial agents were discovered over sixty years ago from a whole cell screening approach against Gram-negative bacteria [6]. Since then, chemistry has expanded quinolone activity to include such agents as delafloxacin, approved in 2017 for treating acute bacterial skin infections caused by the Gram-positive *Staphylococcus aureus*. Some thirteen out of thirty-eight new antibiotics introduced between 2000 and 2019 were quinolones [16-18, 30]. However, safety concerns about quinolone side effects have prompted regulatory recommendations to limit the use of quinolones to patients who do not have other treatment options in both Europe (https://www.ema.europa.eu/en/medicines/human/referrals/quinolone-fluoroquinolone-containing-medicinal-products) and the USA (https://www.fda.gov/drugs/drug-safety-and-availability/fda-drug-safety-communication-fda-advises-restricting-fluoroquino-lone-antibiotic-use-certain).

The determination of the specific DNA sequences cleaved by DNA gyrase or topoisomerase IV [31, 32] was important in determining structures of quinolones in DNA-cleavage complexes. In particular, structural studies showed that quinolone antibiotics stabilize double-stranded DNA-cleavage complexes with the two bacterial topoisomerases, topo IV [33] and DNA gyrase [34, 35], by interacting with ParC or GyrA via a water-metal-ion bridge [12, 36]. Although the original DNA-sequences used in two papers describing structures showing the water metal-ion bridge [33, 35] were defined in 2005 [32] and were initially used in structures with *S. pneumoniae* topo IV [37, 38], they are asymmetric and in high enough resolution structures the DNA was clearly averaged around the twofold axis of the complex [35]. In this paper we used a two-fold symmetric 20-mer DNA duplex to avoid such problems [34, 39]. This 20-mer homoduplex DNA was previously used in determining structures with the progenitor of zoliflodacin, QPT-1 [34].

Herein we describe a 2.8 Å X-ray crystal structure of zoliflodacin in a DNA-cleavage complex with *S. aureus* DNA gyrase. The structure is compared with a structure with the quinolone, moxifloxacin, also in a DNA-cleavage complex with *S. aureus* DNA gyrase. We also show that zoliflodacin has reasonable activity against *A. baumannii* (MICs of 4 *µ*g/*m*L - the A in ESKAPE).

## 2. Results

### 2.1. A 2.8 Å zoliflodacin DNA-cleavage complex with S. aureus DNA gyrase

Crystals of zoliflodacin in a complex with a 20-mer DNA homoduplex (20-447T) and *S. aureus* DNA gyrase were grown by a microbatch crystallization method and a 2.8 Å dataset was collected on beamline I24 at Diamond Light Source (see Materials and Methods for details). The structure was solved from a 2.5 Å QPT-1 complex with the same 20-447T DNA and *S. aureus* DNA gyrase in the same P6_1_ space-group (PDB code: 5CDM; a=b = 93.9 Å, c = 412.5 Å) and refined (see Materials and methods for details – supplementary Figure 1 for electron density). The 20-447T DNA homoduplex contains 18 base-pairs and a G-T mismatch at either end of the DNA.

The structure showed two zoliflodacins binding in the cleaved DNA physically blocking religation (Figure 2). The DNA has been cleaved by and is covalently attached to tyrosine 123A from the GyrA subunit (and to the symmetry related tyrosine 123A’, from the second GyrA subunit in the complex). Catalytic metal ions (normally Mg^2+^ in bacteria) are required for DNA-cleavage, and in our structure, we see two Mn^2+^ ions occupying the ‘B-site’ in the GyrB and GyrB’ subunits [10].

**Figure 2.**
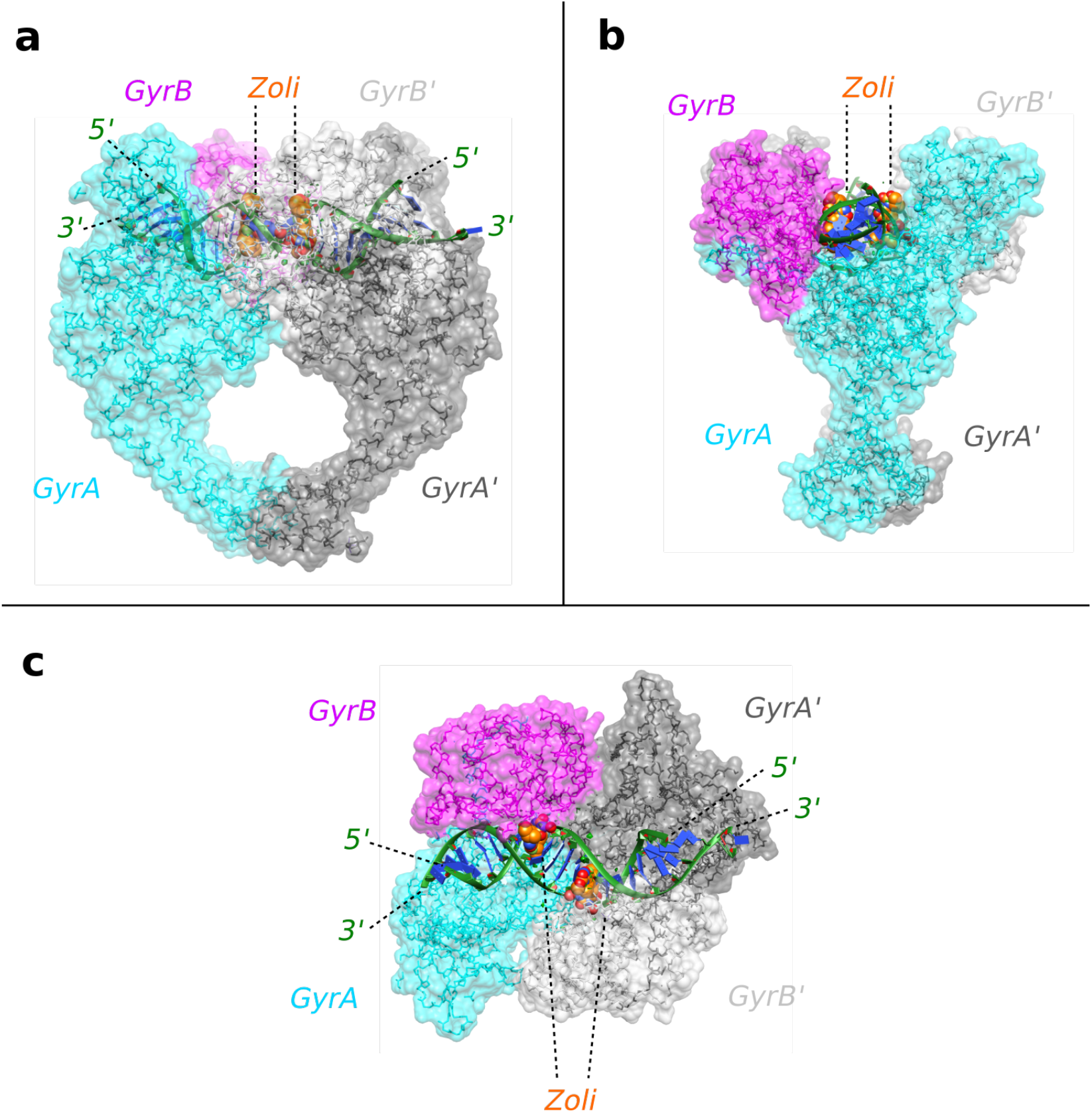
2.8 Å zoliflodacin crystal structure with *S. aureus* DNA gyrase. **(a)** View of 2.8 Å zoliflodacin crystal structure. The DNA (cartoon; green backbone and blue bases) has been cleaved by *S. aureus* gyrase (shown as backbone trace with semi-transparent surface). Tyr 123 (and Tyr 123’) have cleaved the DNA and are covalently attached. The compounds are shown as solid spheres (carbons orange, oxygens red, nitrogens blue). **(b)** An orthogonal (90º) view of the same complex. **(c)** An orthogonal (90º) view looking down the twofold axis of the complex. The two ends of the DNA duplex adopt different conformations due to crystal packing. Figure produced using ChimeraX [40, 41].

### 2.2. Zoliflodacin interacts with GyrB, whereas moxifloxacin interacts with GyrA

Figure 3 compares the binding sites of compounds in our 2.8 Å zoliflodacin structure with a 2.95 Å *S. aureus* DNA gyrase DNA-cleavage complex with the widely used quinolone antibiotic, moxifloxacin (PDB code: 5CDQ [34]). Figure 3a shows the binding mode of zoliflodacin with the pyrimidinetrione (or barbituric acid moiety) of the compound making direct interactions with GyrB. In particular, the terminal oxygen of the pyrimidinetrione makes a hydrogen bond with the main-chain NH of aspartic acid B437.

**Figure 3.**
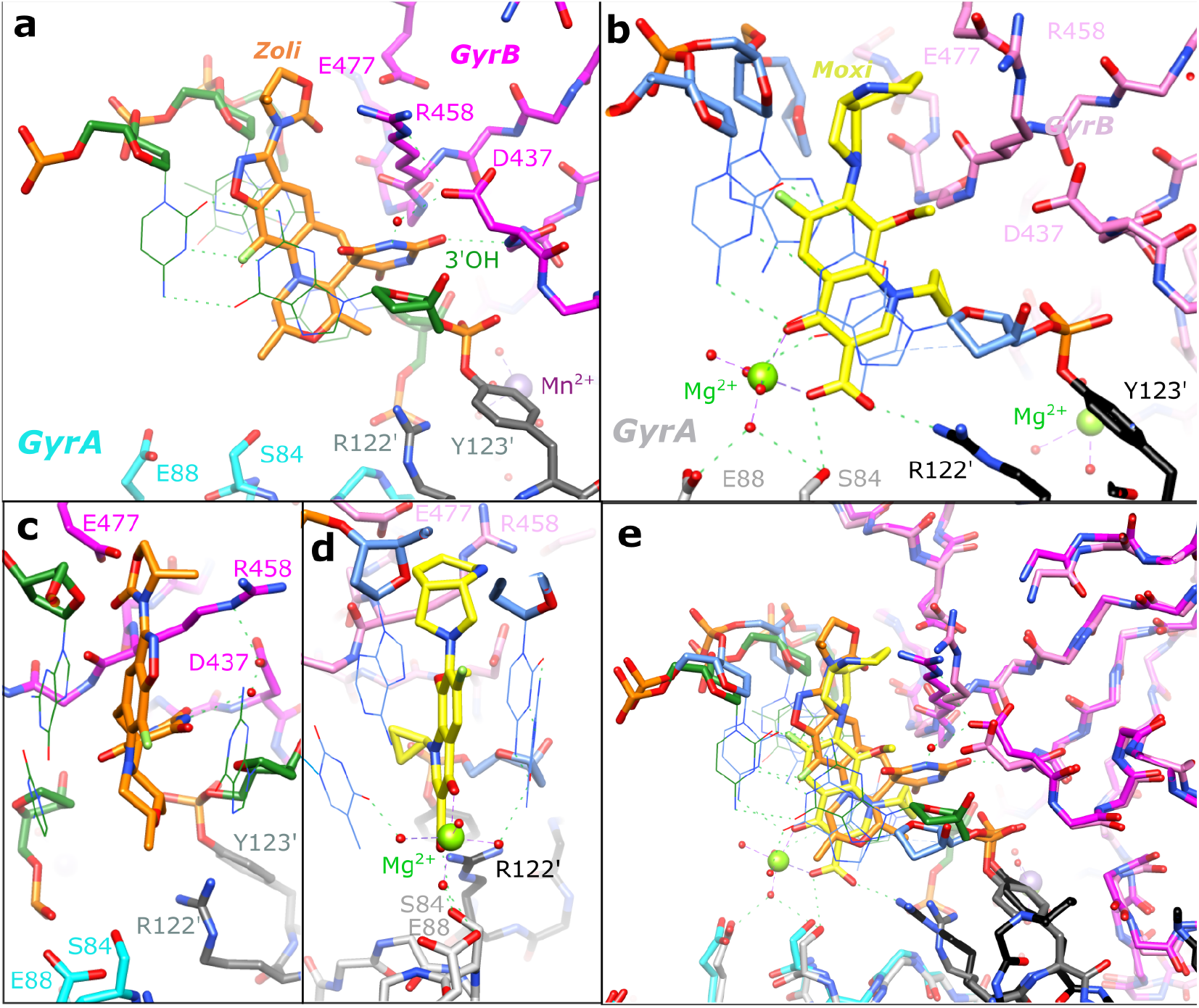
Equivalent views of zoliflodacin and moxifloxacin in *S. aureus* DNA gyrase DNA-cleavage complexes. (**a**) A close-up view of a zoliflodacin binding site in the 2.8 Å structure. The pyrimidinetrione moiety of zoliflodacin interacts directly and indirectly (via a water) with Asp437 of GyrB (dotted green lines), in a similar manner to that described in the 2.5 Å QPT-1 structure. Y123’ has cleaved the DNA and forms a ‘phosphotyrosine’ type linkage with the cleaved DNA. The DNA-backbone is shown in fatter ‘stick’ representation with the bases drawn in thinner ‘line’ (base-pair H-bonds only shown for the +1, +4 base-pair). (**b**) In the 2.95 Å moxifloxacin (yellow carbons) structure the quinolone bound Mg2+ ion (green sphere) and coordinating waters (red spheres) make hydrogen bonds (dotted red lines) to S84, E88 and the bases either side of the DNA-cleavage site (at the +1 and -1 positions - see panel d). (**c** + **d**) Orthogonal (90º) views of the compound binding sites in zoliflodacin structure (c) and moxifloxacin structure (d). (**e)** Superposition of **a** and **b**. Figure produced using ChimeraX [40, 41].

This contrasts with moxifloxacin where the compound (Figure 3b and 3e) interacts with S84A and E88A from the GyrA subunit via the now well characterized water-metal ion (Mg^2+^) bridge [12, 33-35, 42, 43]. The lack of interactions with GyrA and the interactions with GyrB account for the much of the activity of zoliflodacin against quinolone resistant strains of bacteria (e.g. Table 4 in [3]; target mediated resistance is common in quinolone resistant bacteria [11]). The interactions of the quinolones with the GyrA (or ParC) subunit via the flexible water-metal ion bridge may account for some of the specificity of quinolones for DNA gyrase and topo IV (see sequence alignment in Figure 4) over the two human type IIA topoisomerases, Top2α and Top2β.

**Figure 4.**
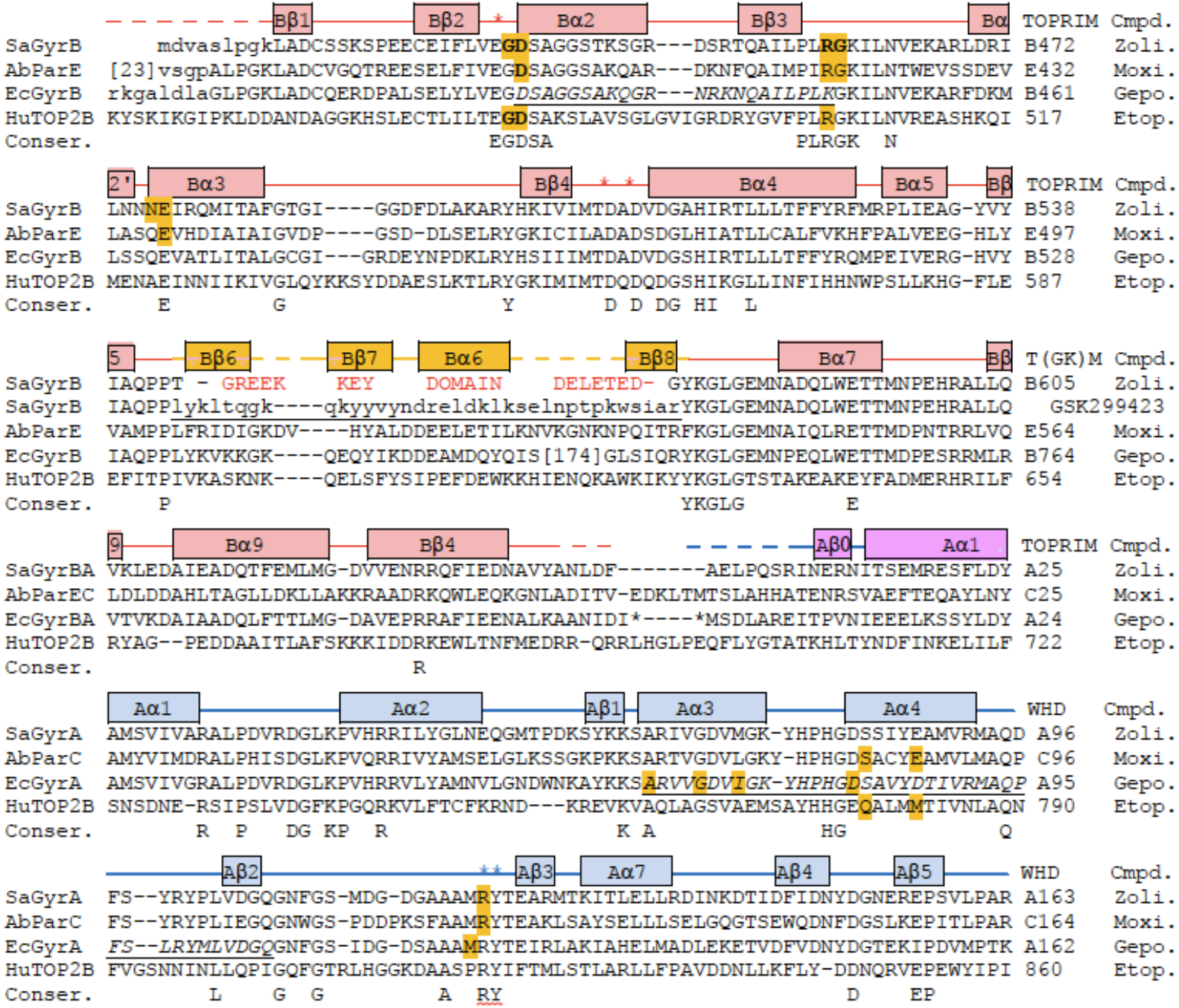
Sequence alignment highlighting residues contacting zoliflodacin, moxifloxacin, gepotidacin or etoposide. An alignment of resides in TOPRIM, Greek Key and WHD (N-terminal part of) domains of: *S. aureus* GyrB/GyrA (SaGyrBA), *A. baumannii* ParE/ParC (AbParEC), *E. coli*. GyrB/GyrA (EcGyrBA) and human Top2β (HuTOP2B). Residues in ‘Conser.’ are conserved in eight aligned sequences in supplementary Figure 5 from Chan *et al*., 2015 [34] (five bacterial topo2As and the three eucaryotic topo2As; note the central R in PLRGK can be conservatively varied to K). Amino acids are highlighted in sequences if they contact (< 3.8 Å) compounds in *S. aureus* DNA gyrase structures with compounds. Contacts mapped onto the *S. aureus* DNA gyrase sequence are Zoli. = contacts in the 2.8 Å zoliflodacin structure (PDB code: 8BP2). The contacts mapped onto *A. baumannii* topo IV (ParE/ParC) sequence are Moxi. = contacts from the 2.95 Å moxifloxacin complex (PDB code: 5CDQ: note for moxifloxacin the Mg2+ ion and waters of the water metal-ion bridge are taken as part of the compound. Contacts in a 3.25 Å *A. Baumannii* topo IV moxifloxacin complex structure, 2XKK, are nearly identical to those shown). The contacts mapped onto *E. coli* Gyrase sequence are Gepo. = from the 2.37 Å gepotidacin with uncleaved DNA (PDB code: 6QTP; note contacts in the 2.31 Å structure 6QTK, and in the full-length *E. Coli* CryoEM gepotidacin structures. e.g PDB code: 6RKS are very similar). The contacts mapped onto the HuTOP2B structure are from the 2.16 Å etoposide complex with human Top2β (PDB code: 3QX3 - but are similar in *S. aureus* etoposide crystal structures: 5CDP and 5CDN). The secondary structural elements in a 3.5 Å *S. aureus* gyrase complex with DNA and GSK299423 (2XCR) are shown above the alignment - note deletion of the greek key domain gave a GSK299423 structure at 2.1 Å (2XCS). The * above the sequence alignment indicates positions of catalytic residues (Glu B435, AspB508, Asp B510, Arg A122, Tyr A123 in *S. aureus* DNA gyrase). The <u>quinolone resistance-determining region (QRDR</u>), defined as 426-447 in *E. coli* GyrB 67-106 in *E. coli* GyrA is <u>underlined in italics</u> on the EcGyrB/A sequences [11].

However, the residues on GyrB which partly form the DNA gyrase-zoliflodacin binding interactions, are conserved not only in the bacterial type IIA topoisomerases but also in the human enzymes. As shown in the sequence alignment in Figure 4, and in Figure 3a, zoliflodacin recognizes and interacts with the GD from the conserved E<u>GD</u>SA motif and the RG from the PL<u>RG</u>K. While E435 at the start of the EGDSA motif is a catalytic residue the other residues from the EGDSA are not catalytic, neither are the PLRGK motif residues. In QPT-1 only the GD and RG residues from GyrB contacts the compound The specificity of spiropyrimidinetriones, such as zoliflodacin, towards bacterial type IIA topoisomerases such as human topoisomerases is believed to be because such compounds can be squeezed out of the pocket when the DNA-gate closes in human topoisomerases [34]. Conformational flexibility in spiropyrimidinetrione ligands, such as QPT1 and zoliflodacin, may be important in allowing ligands to maintain favorable interactions within the binding sites as the DNA wriggles the protein [34, 44, 45]. In addition to the multiple tautomeric forms that the pyrimidinetrione moiety can adopt (only one of which is chemically called a ‘pyrimidinetrione’), and the conformational flexibility of the anilino-nitrogen [34], the methyl-oxazolidine-2-one may also be able to adopt more than one conformation. Multiple high-resolution structures will be required to fully discern how the compound wriggles (when its binding pocket changes shape) as the enzyme is moved around by its substrate DNA [45]. However, from this initial 2.8 Å zoliflodacin structure it is clear that the major protein interactions made by zoliflodacin are clearly with the GD and the RG from the highly conserved E**<u>GD</u>**SA and PL**<u>RG</u>**K motifs (Figures 3 and 4).

In the 2.1 Å crystal structure of the NBTI GSK299423 with the *S. aureus* gyrase^CORE^ and DNA (PDB code: 2XCS; [13]), a Y123F mutant was used so the DNA could not be cleaved. In this 2.1 Å GSK299423 structure the +1:+4 base-pair (Figure 1c) occupies a similar space to the inhibitors in the zoliflodacin and moxifloxacin structures. Some reasons for the conservation of the EGDSA and PLRGK motifs (Figure 4) may be discerned from this 2.1 Å structure. While the side-chain of E435 (the first residue of the EGDSA motif) coordinates the catalytic metal (at the ‘A’ position - poised to cleave the DNA) both the main-chain NH and side-chain hydroxyl of serine 438 are within hydrogen-bonding distance of the phosphate between nucleotides 1 and 2. The main-chain C=O of Arg 458 and the main-chain NH of Lys 460 (from the PLRGK motif), accept and donate hydrogen bonds to the -1 guanine base helping to hold it firmly in place. NBTIs can stabilize complexes with one strand cleaved or with no DNA-cleavage [13, 24, 46], however experimental nucleotide preferences for NBTI-cleavage have not yet, to the best of our knowledge, been determined [47].

### 2.3 Target mediated resistance to Zoliflodacin in Neisseria gonorrhoeae

The binding of zoliflodacin to the conserved motifs on GyrB correlates well with the low prevalence of target-mediated resistance; only one of some 12,493 *N. gonorrhoeae* genomes from the PathogenWatch database has a predicted first level resistance mutation [48]. Assessing the probability of developing resistance is an important step in development of any new antibiotic. The development of zoliflodacin (AZD0914) for gonorrhea followed from a 2015 paper assessing the likelihood of developing resistance in *N. gonorrhoeae* [49]. This paper showed that higher MICs (resistance) was associated with target mutations in three amino acids in *N. gonorrhoeae* DNA gyrase, namely GyrB: D429N, K450T or S467N [49]. These mutations were identified by in vitro selection of resistance and can give a 4-fold to 16-fold increase in the MIC of zoliflodacin [49, 50]. Interestingly these *N. gonorrhoeae* GyrB mutations correspond to D437, R458 and N475 in *S. aureus* DNA gyrase. The D429N mutation is associated with slower growth of bacteria [51]. All three regions are close to the compound (see Figure 3a). In the D437N mutant (*S. aureus* DNA gyrase) the asparagine side chain may have its NH_2_ group pointing towards the compound (because if the sidechain was in the opposite orientation the hydrogens on the NH_2_ would clash with hydrogens on proline 56 - the P in PLRGK).

Zoliflodacin has an extra methyl-oxazolidine-2-one ring, which QPT-1 does not possess (Figure 1a, b) and this extra ring makes van der Waals contacts with residues N476 and E477. While there is clear electron density for the both the additional fluorine and the extra ring (which are not in the QPT-1 structure - Figure 5), the 2.8 Å electron density map is not able to clearly define all water structure or totally unambiguously define the orientation of the extra five-membered ring (see supplementary Figure 1). N475 is equivalent to the third mutated residue in *N. gonorrhoeae* GyrB, Ser 467 [49]. Mutation of this residue, which is adjacent to residues contacting the compound, presumably effects their conformations. A similar effect is perhaps seen in the *S. aureus* ParC V67A, found in a strain of *S. aureus* resistant to gepotidacin [52]. In high resolution *S. aureus* DNA gyrase NBTI crystal structures three residues (**<u>A</u>, <u>G</u>** and **<u>M</u>**) from the GyrA motif 68-**<u>A</u>**RIV**<u>G</u>**DV**<u>M</u>**-75 are within Van Der Waals distance of compounds [13, 22, 24]. ParC V67A is the first V in the equivalent ParC sequence <u>A</u>KTV<u>G</u>DVI, i.e Val 67 is adjacent to an amino acid making direct Van Der Waals contacts with compounds.

**Figure 5.**
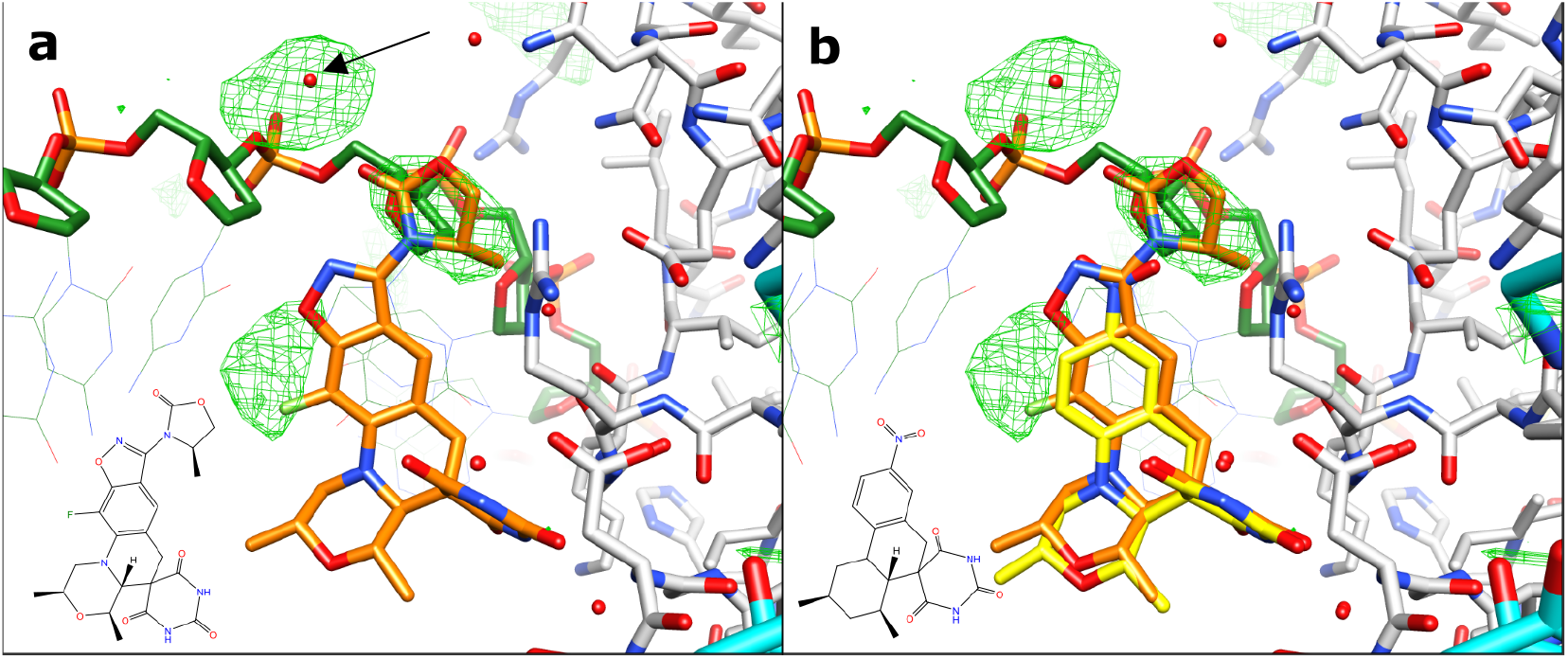
Difference map density from refined QPT-1 structure against zoliflodacin data. **(a)** An Fo-Fc map from refined QPT-1 coordinates is contoured at 3 σ shows extra features in zoliflodacin structure not in QPT-1 starting coordinates. The arrow points to extra density modelled as a water – which could be a metal ion (it is close to the oxygen of a phosphate from the DNA backbone and also an oxygen from the methyl-oxazolidine-2-one) **(b)** refined QPT-1 coordinates are also shown with yellow carbons.

### 2.5 Improved activity of zoliflodacin against A. baumannii compared to QPT-1

The MIC of zoliflodacin against two carbapenem resistant outbreak strains of *Acinetobacter baumannii* [53] was determined (see Materials and methods for details) as 4 *µ*g/*m*L (Table 1). This suggests that, although its activity has been optimized against other Gram-negative bacteria, that the potency of zoliflodacin against *A. baumannii* is better than that of QPT-1 from which it was developed (the activity of QPT-1 against *A. baumannii* is from Supplementary Table 4 in Chan et al., (2015); but note QPT-1 is considerably more active against an efflux knock-out strain: *A. baumannii* BM4454 (βadeABC βadeIJK) [54]).

**Table 1.** MIC of zoliflodacin for *A. baumannii*.

| 283 | Compound | Species | MIC (µg/mL) |
| --- | --- | --- | --- |
|  | Zoliflodacin | *A. baumannii* BCC 807 (UK OXA-23 clone) | 4 |
|  | Zoliflodacin | *A. baumannii* BCC 810 (South East OXA-23 clone) | 4 |
|  | QPT-1 | *A. baumannii* BM4454 | 16 |
|  | QPT-1 | *A. baumannii* BM4454 (ΔadeABC ΔadeIJK)* | 0.125 |

\* *A. baumannii* BM4454 (adeABC and adeIJK) is equivalent to BM4652 [54].

## 3. Discussion

Some Gram-negative bacteria are difficult to kill with antibiotics. Not only do they have two cell walls but they also have export pumps that can rapidly pump antibiotics out of the bacteria [54]. Such bacterial export pumps can play a role in antimicrobial resistance [55]. There is much interest in compounds that can inhibit antibiotic efflux pumps [56], as there is clearly a potential for combination therapies. If the MICs of a compound such as zoliflodacin could be lowered, the dose might be lowered, and the therapeutic window increased.

However, the success of whole cell screening, including early counter-screening of human cells for safety seems to have been effective in discovering two new classes of Gram-negative targeting antibiotics [5, 13]. A similar approach, but starting with a natural product, has recently lead to the discovery of evybactin a new class on *M. tuberculosis* DNA gyrase targeting compound [57]; this compound appears to work in a similar manner to the thiophene inhibitors [58] compounds that allosterically stabilize DNA-cleavage complexes [59] by binding to a ‘third-site’; a hinge pocket [10].

Interestingly two of the mutations in *N. gonorrhoeae* GyrB that give rise to resistance to zoliflodacin (AZD0914) are Asp429Asn and Lys450Thr, correspond in *S. aureus* crystal structures to residues involved in making up the binding pocket of the compound (Figure 3). Namely Asp 437 (=Asp429) is the D from the conserved EGDSA motif and Arg 458 (=Lys450) is from the conserved PLRGK motif.

In a previous paper describing crystals structures of QPT-1, moxifloxacin and etoposide in DNA-cleavage complexes with *S. aureus* DNA gyrase [34], the DNA-gate of DNA-gyrase was proposed to act like a pair of swing doors, through which the T-segment could be pushed (Figure 1f) but that would then swing close. Such a model might partly account for why in *N. gonorrhoeae* mutations are only seen in GyrB not in ParE [49]. The swinging closed of the DNA-gate in DNA gyrase might be predicted to give slower ‘off’ rates for zoliflodacin compared to topoisomerase IV; note it was also proposed that zoliflodacin would be squeezed out of a slightly larger equivalent pocket in human topo2s [34]. Much work remains to be done; for example, one current model suggests that before the C- (or exit) gate can be opened the small greek key domain senses the presence of the T-DNA segment (once it has passed through the G-gate) and then moves the catalytic metal away from the active site (see supplementary discussion and supplementary Figures 12 and 13 in [34]). This model allows the DNA to be religated by the lysine residue from the highly conserved Y**<u>K</u>**GLG motif at the C-terminus of the greek key domain (see Figure 4), while not allowing DNA-cleavage by the catalytic metal when the exit gate is opened, and not allowing exit-gate opening while the gate-DNA is cleaved. In this model this is a ‘safety feature’ of type IIA topoisomerases - allowing DNA religation by the Y**<u>K</u>**GLG lysine but inhibiting DNA-cleavage by the catalytic metal. Interestingly it has also been shown that DNA-gyrase can catalyze supercoiling by introducing a single nick in the DNA [60], perhaps this mechanism is also a safe way of introducing negative supercoils into DNA without opening the C-gate.

Safety and the size of the therapeutic window are clearly important in antibacterial drug discovery. It will be interesting to see if the new spiropyrimidinetrione class of compounds, such as zoliflodacin, can be developed to be safer and more efficacious medicines, with less of a tendency for target-mediated antibiotic resistance, than the quinolones.

## 4. Materials and Methods

### 4.1 Protein purification and crystallization of a zoliflodacin DNA-cleavage complex

The *S. aureus* DNA gyrase fusion truncate GyrB27:A56 (GKdel) (M_w_ 78,020) was expressed in *E. coli* and purified based on the procedure of Bax *et al*., [13] modified as described [25]. The purified protein (at 10 *m*g/*m*L = 0.128 *m*M) was in 20 *m*M HEPES pH 7.0, 5 *m*M MnCl_2_ and 100 *m*M NaSO_4_. The DNA oligonucleotide used in crystallizations, 20-447T, was custom ordered from Eurogentec (Seraing, Belgium). Received in lyophilized form, the DNA was resuspended in nuclease-free water and annealed from 86 to 21°C over 45 minutes to give the duplex DNA at a concentration of 2 *m*M. Zoliflodacin was purchased from MedChemExpress (New Jersey, USA) as a solid and was dissolved in 100% DMSO forming a 100 *m*M stock solution.

Crystallization complexes were formed by mixing protein, HEPES buffer, DNA and compound and incubating on ice for 1 hour 15 minutes. Crystals of *S. aureus* GyrB27:A56 (GKdel)-zoliflodacin-20-447T were grown by the microbatch under oil method [39], with streak seeding being implemented for subsequent plates after the first plate gave crystals. Following established protocols, a crystallization screen consisting of Bis-Tris buffer pH 6.3 - 6.0 (90, 150 *m*M) and PEG 5*k*MME (13 – 7%) was used. For a single drop, 1 *µ*L of complex mixture was mixed with 1 *µ*L of crystallization buffer in a 72-well Terasaki microbatch plate, prior to covering with paraffin oil. The plates were incubated at 20°C and crystal growth was observed between 5 and 30 days. Seed solution was prepared by crushing several previously grown hexagonal rod-shaped *S. aureus* GyrB27:A56 (GKdel)-zoliflodacin-20-447T crystals in 20 *µ*L of crystallization buffer. The crystal which gave a 2.8 Å dataset was grown in a crystallization plate where 1 *µ*L complex mixture (0.066 *m*M GyrB27:A56 dimer, 0.171 *m*M 20-447T DNA duplex, 5.714 *m*M zoliflodacin, 2.571 *m*M MnCl_2_ and 342.9 *m*M HEPES pH 7.2) was mixed with 1 *µ*L crystallization buffer (90 *m*M Bis-Tris pH 6.3, 9% PEG 5 *k*MME). A large single crystal (Figure 1) was transferred to a cryobuffer (15% glycerol, 19% PEG 5*k*MME, 1 *m*M zoliflodacin, 5% DMSO, 81 *m*M Bis-Tris pH 6.3) before flash-cooling in liquid nitrogen for data collection.

### 4.2 Data collection, structure determination and refinement

Data were collected (3600 × 0.lº degree images) on beamline I24 at Diamond Light Source. Data were processed and merged with dials **[61-63]** as shown in Table 2. A low-resolution cutoff of 25 Å was applied when manually reprocessing the data with dials to avoid problems with the backstop shadow. The high-resolution cutoff was determined by having a **CC**_**1/2**_ **> 0.30 [64]**. The structure was refined starting from the 2.5 Å complex with the same DNA and the related compound QPT-1 (PDB code: 5CDM) [34, 39]. The data, which are not twinned and are in space-group P6_1_, were reindexed (H=k, K=h, L=-l) to be in the same hand and on the same origin as other liganded structures in the same space-group (e.g PDB codes: 2XCS, 4BUL, 5IWI, 5IWM, 5NPP, 6QTK, 6QX1 and 5CDM). Initial rigid body refinement of 5cdm-BA-x.pdb (P6_1_ cell: a=b=93.88Å, c=412.48Å), reduced the R-factor (R-free) from 0.3900 (0.3899) to 0.2684 (0.2743). Further refinement with phenix.refine [65, 66] and refmac [67, 68] gave the final structure (Table 3), which had reasonable geometry. Restraints for zoliflodacin were generated in Acedrg [69]. As we are interested in structures with ligands/inhibitors we use the standard BA-x numbering scheme throughout this paper [10]. This means zoliflodacin inhibitors in sites 1 and 1’ have CHAINID I, and residue numbers 1 and 201 (see Figure 3 in ref. [10]). Electron density maps for the inhibitors are shown in supplementary Figure 1. The water structure near the inhibitors was based on that in the 2.5 Å structure with QPT-1 (PDB code: 5CDM). Water and glycerol structure was based on electron density maps and higher resolution structures (the 1.98 Å *S. aureus* complex PDB code: 5IWI, which contains over 940 waters, was superposed).

**Table 2.**
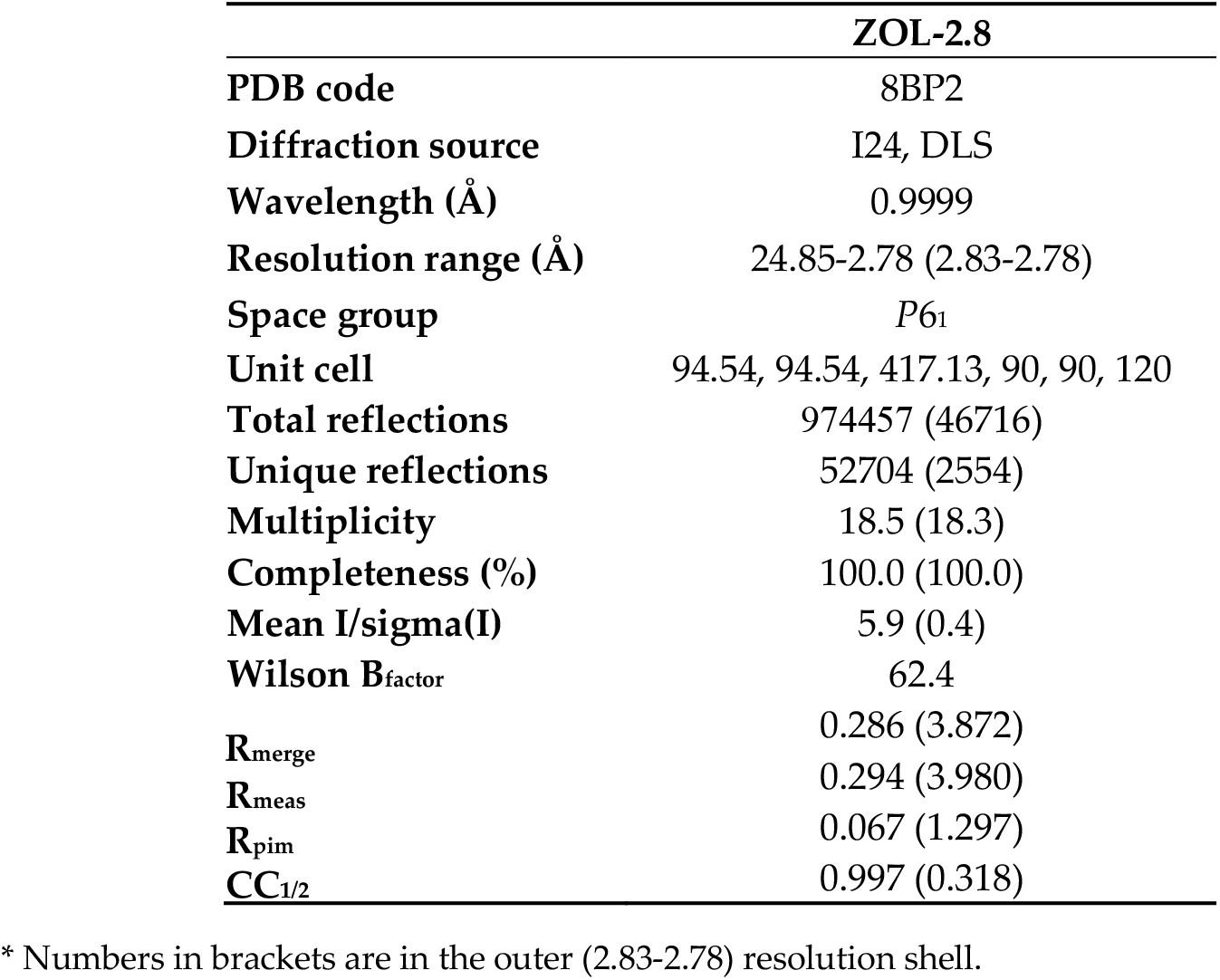
Data collection statistics.

**Table 3.** Refinement statistics.

|  | **ZOL-2.8** |
| --- | --- |
| **PDB code** | 8BP2 |
| **Resolution range (Å)** | 24.85-2.80 (2.87-2.80) |
| **Completeness (%)** | 99.41 (94.60) |
| **No. of reflections, working set** | 48785 (3421) |
| **No. of reflections, test set** | 2538 (185) |
| **Final Rcryst** | 0.1957 (0.372) |
| **Final Rfree** | 0.2375 (0.374) |
| **Cruickshank DPI (Å)\*** | 0.325 |
| **No. of non-H atoms (total)** | 11713 |
| **Protein** | 10574 |
| **DNA** | 801 |
| **Zoliflodacin** | 70 |
| **Other ligands (Mn, glycerol etc.)** | 42 |
| **Waters** | 226 |
| **RMS deviations** |  |
| **Bonds (Å)** | 0.009 |
| **Angles (°)** | 1.569 |
| **Average B factors (Å<sup>2</sup>)** |  |
| **Protein** | 96.024 |
| **DNA** | 85.103 |
| **Zoliflodacin** | 84.491 |
| **Other ligands (Mn, glycerol etc.)** | 88.911 |
| **Waters\*\*** | 75.396 |
| **Ramachandran plot** |  |
| **Favored regions** | 96% |
| **Additionally allowed** | 4% |
| **Outliers** | 0% |

\* The Cruickshank DPI (Å) was calculated by the Online\_DPI server [70](Kumar et al., 2015).
\*\* Waters were placed where there are waters in higher resolution structures (e.g. 5CDM and 5IWI).

At each DNA-cleavage site a single catalytic Mn^2+^ ion is seen at the B-site [10]. Electron density on His C 391, was interpreted as being due to a Mn^2+^ ion coordinated by a Bis-Tris buffer molecule, which mediates a crystal contact with one end of the DNA. This interpretation of the electron density was confirmed by re-refining the original 2.1 Å structure of GSK299423 with the *S. aureus* gyrase^CORE^ structure [13]; originally this electron density had been misinterpreted as being due to DNA. The new interpretation explains why both Bis-Tris and Mn^2+^ ions are needed in the crystallization buffer, when growing P6_1_ crystals of *S. aureus* gyrase^CORE^ with ligands.

### 4.3 Structural analysis

Van der Waals contacts with ligands (defined as 3.8 Å or less) were calculated with contact from the CCP4 suite of programs [71]. Structures were superposed using coot [72] or with limited sets of defined Cαs using LsqKab from CCP4 suite [71].

### 4.4 Minimum Inhibitory Concentration Assay

The MIC of zoliflodacin against two carbapenem resistant outbreak strains of *A. baumannii* (BCC 807, BCC 810) [53] was determined using the modified broth microdilution reference method ISO 20776-1:2019 [73] as recommended by the EUCAST (European Committee on Antimicrobial Susceptibility Testing) [74]. The concentration range tested was between 40 – 0.313 *µ*g/*m*L in two-fold serial dilutions.

## Supplementary Materials

The following supporting information can be downloaded at: www.mdpi.com/xxx/s1, Figure S1: Difference density (Fo-Fc) for the two zoliflodacins.

## Author Contributions

“Conceptualization, B.D.B. and H.M.; protein purification J.H. and N.P.B; complex prepara-tion and crystallization H.M and M.L.-L.; data-collection and processing, H.M. and A.W.; structure determination and analysis H.M and B.D.B.; microbiology and MIC determination N.H. and E.M.; writing—original draft preparation, B.D.B. and H.M.; writing—review and editing, all authors. All authors have read and agreed to the published version of the manuscript.”

## Data Availability Statement

The 2.8 Å zoliflodacin structure has been deposited with the protein databank (the PDB) with the code: 8BP2. The PDB nomenclature for compounds and metal ions is inconsistent between different structures. So sensibly named PDB files, following BA-X nomenclature [10] are made available from a table of structures on the ‘Research’ tab on Ben Bax’s website at Cardiff (https://www.cardiff.ac.uk/people/view/1141625-bax-ben).

## Acknowledgments

We thank Professor Simon Ward and the Medicines Discovery Institute (MDI) in Cardiff for supporting this work. We thank Diamond Light Source (DLS) for beamtime and general support. HM is a joint student between the MDI and DLS.

